# Centrifuger: lossless compression of microbial genomes for efficient and accurate metagenomic sequence classification

**DOI:** 10.1101/2023.11.15.567129

**Authors:** Li Song, Ben Langmead

## Abstract

Centrifuger is an efficient taxonomic classification method that compares sequencing reads against a microbial genome database. In Centrifuger, the Burrows-Wheeler transformed genome sequences are losslessly compressed using a novel scheme called run-block compression. Run-block compression achieves sublinear space complexity and is effective at compressing diverse microbial databases like RefSeq while supporting fast rank queries. Combining this compression method with other strategies for compacting the Ferragina-Manzini (FM) index, Centrifuger reduces the memory footprint by half compared to other FM-index-based approaches. Furthermore, the lossless compression and the unconstrained match length help Centrifuger achieve greater accuracy than competing methods at lower taxonomic levels.

## Background

Metagenomic sequencing enables comprehensive profiling of microbiomes in a sample and has been widely applied to study natural environments [1,2], infectious diseases [3], allergies [4] and cancers [5]. Taxonomic classification labels each sequencing read with taxonomy IDs representing its most likely taxon of origin. This has become an important step in translating raw sequencing data into meaningful microbiome profiles [6]. Classification is usually conducted by comparing the read sequence to all the sequences in a database of microbial reference genomes, such as RefSeq [7], GeneBank [8], or GTDB [9]. The growth of available microbial reference genomes creates a strong need for memory-efficient structures. Many methods turn to lossy representations of the database. For example, Kraken2 [10] reduces the space by storing minimizers [11] instead of all the k-mers as in Kraken [12]. Other approaches, such as MetaPhlAn [13,14] and CLARK [15], build the database out of only a selected subset of sequences, i.e., marker genes or discriminative k-mers. Ganon [16] and KMCP [17] utilize probabilistic data structures that discard k-mer identity but support checking k-mer presence with false positive probability. While these strategies reduce the memory requirement, they lose valuable sequence information, which may lower accuracy when classifying read to lower taxonomic levels. We previously developed the taxonomic classification method Centrifuge [18] that used the memory-efficient Burrows-Wheeler transformed (BWT) sequence [19] and the Ferragina-Manzini (FM) index [20]. Centrifuge searches for semi-maximal matches with no length constraints, avoiding the decreased taxonomic specificity of k-mers when the genome database is large [21]. However, the FM-index grows linearly with the database size and the lossy compression strategy proposed in Centrifuge is not scalable, making Centrifuge less usable in the context of large and growing genome databases.

Related genomes share similar sequences, giving genome databases a degree of repetitiveness. Lempel-Ziv family indexes [22], context-free grammars [23], and run-length compressed BWT indexes (RLBWT) [24] exploit this repetitiveness to reduce index size losslessly while supporting efficient search queries. For example, r-index [25] builds upon the RLBWT and fits in O(r) words, where r is the number of runs in the BWT sequence. The FM Index, by contrast, usually requires O(n) words.

While O(r)-space methods achieve good compression for highly repetitive sequences such as collections of human genomes, microbial genomes are more diverse. Applying these compact representations may take more space than the uncompressed wavelet tree [26]. Therefore, we designed two compact data structures, called run-block compressed BWT (RBBWT) and hybrid run-length compressed BWT, to effectively compress the BWT sequence for the intermediate level of repetitiveness characteristic of microbial genome databases. RBBWT achieved the best overall performance in both time and space efficiency when compared to other compression methods. Inspired by the observation, we developed the software tool Centrifuger (Centrifuge with RBBWT), which rapidly assigned the taxonomy IDs for a sequencing read while consuming half the memory of a conventional FM-index.

## Results

### Method overview

Centrifuger assigns a taxonomic ID to each input read or read pair by searching against a losslessly compressed FM-index built from a microbial genome database (Figure 1). The FM-index contains two memory-consuming components, the data structure supporting rank queries over the BWT sequence, and the subsampled suffix array. We propose a novel compact structure, the run-block compressed sequence, to reduce the size needed to store the BWT sequence. For the sampled suffix array, we save space by storing only the sequence ID for each sampled position, omitting information about the offset within the genome. The classification algorithm scans the read twice, once for the original sequence and once for the reverse-complement sequence. Each scan looks for semi-maximal matches, repeatedly extending the match with the backward search, then skipping the base immediately after the point where the backward search terminates. For each match, Centrifuger retrieves the sequence IDs associated with entries in the matching BWT range. Centrifuger adds a score, which is a quadratic function of the match length, for each retrieved sequence ID. Aber all the valid matches are processed, the highest-scoring taxonomy IDs translated from the sequence IDs are reported as the classification result. When the number of reported IDs exceeds the user-specified threshold (default 1), Centrifuger reduces the number to within the threshold by promoting some taxonomy IDs to their lowest common ancestors (LCAs) in the taxonomy tree.

**Figure 1:**
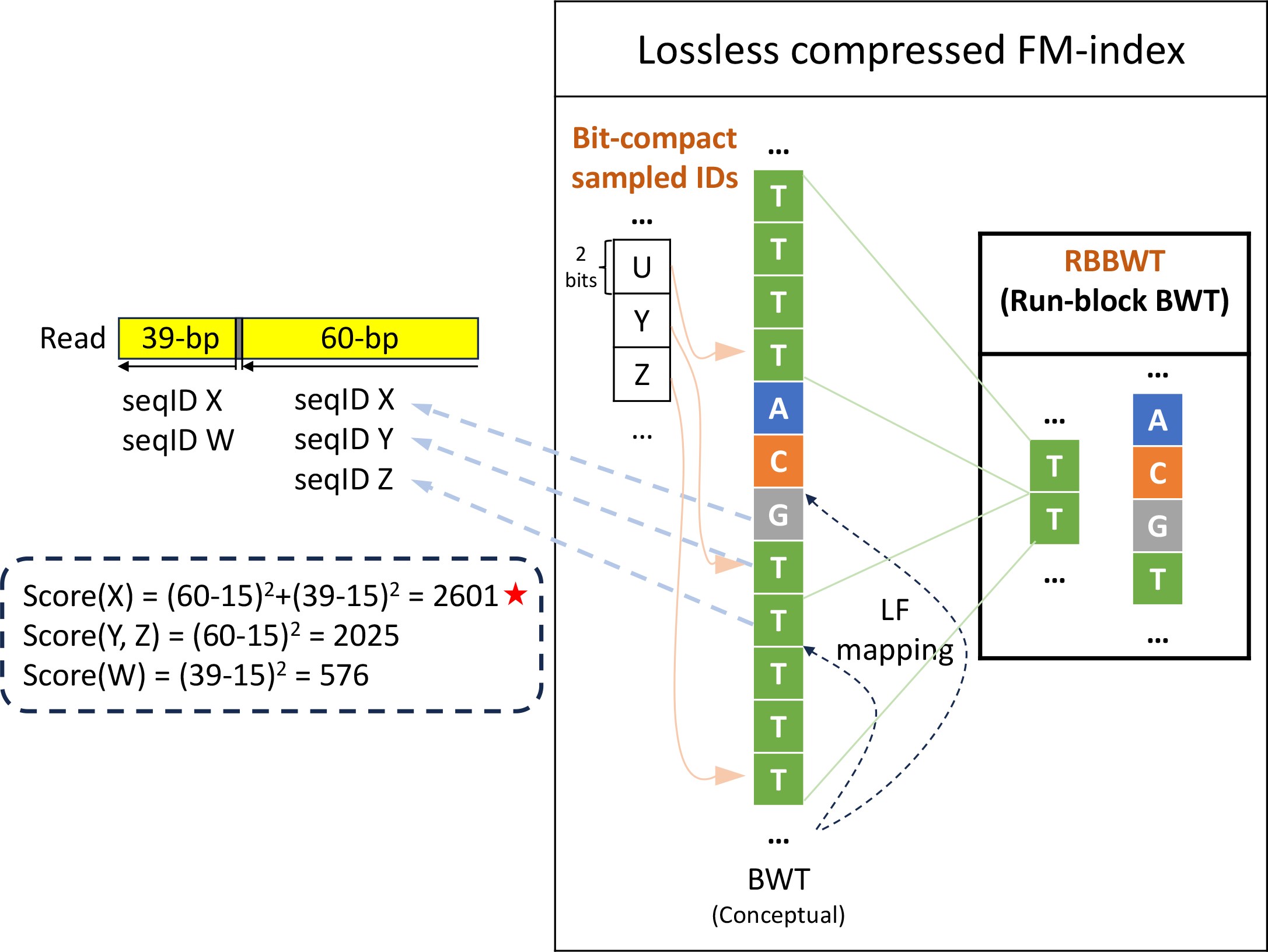
Overview of Centrifuger

### The computational efficiency of run-block compression

Run-block compressed sequence is a novel compact data structure that achieves sublinear space usage both in theory 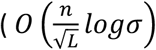 bits. L: average run length, i.e.,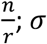 alphabet size) and practice. Centrifuger applies run-block compression to reduce the size of the BWT sequence, yielding the RBBWT: Run-Block compressed BWT. Rank queries on the RBBWT, which form the basis for LF-mapping in the backward search, are also highly efficient, having a time complexity of *O*(*log*σ) (More information in Method section).

We compared RBBWT with three other representations of BWT sequences: the standard wavelet tree, the RLBWT as implemented in the r-index package [27] using sdsl library [28], and the hybrid run-length compression (Method). We measured the change in space usage when adding non-plasmid sequences from the species *Escherichia fergusonii* (taxonomy ID 564) to the structure. While the wavelet tree grew linearly as more genomes were added, RBBWT, RLBWT and its hybrid version grew more slowly (Figure 2A left). When there was little repetitiveness in the genomes, RBBWT and hybrid run-length compression took almost the same amount of space as the wavelet tree. From another perspective, when the average run length of the BWT increased, the number of bits to represent a nucleotide in the wavelet tree remained constant (0.31 bits/bp in our implementation), and the other three compression methods needed fewer bits (Figure 2A right). RBBWT consumed the least or similar space compared with the runlength-based compression methods when L was less than 10. When the BWT was constructed from all the genomes under taxonomy ID 564 with L equaling 18.8, RBBWT was still small, consuming 57.8% less space than the wavelet tree and 29.8% more space than RLBWT.

**Figure 2.**
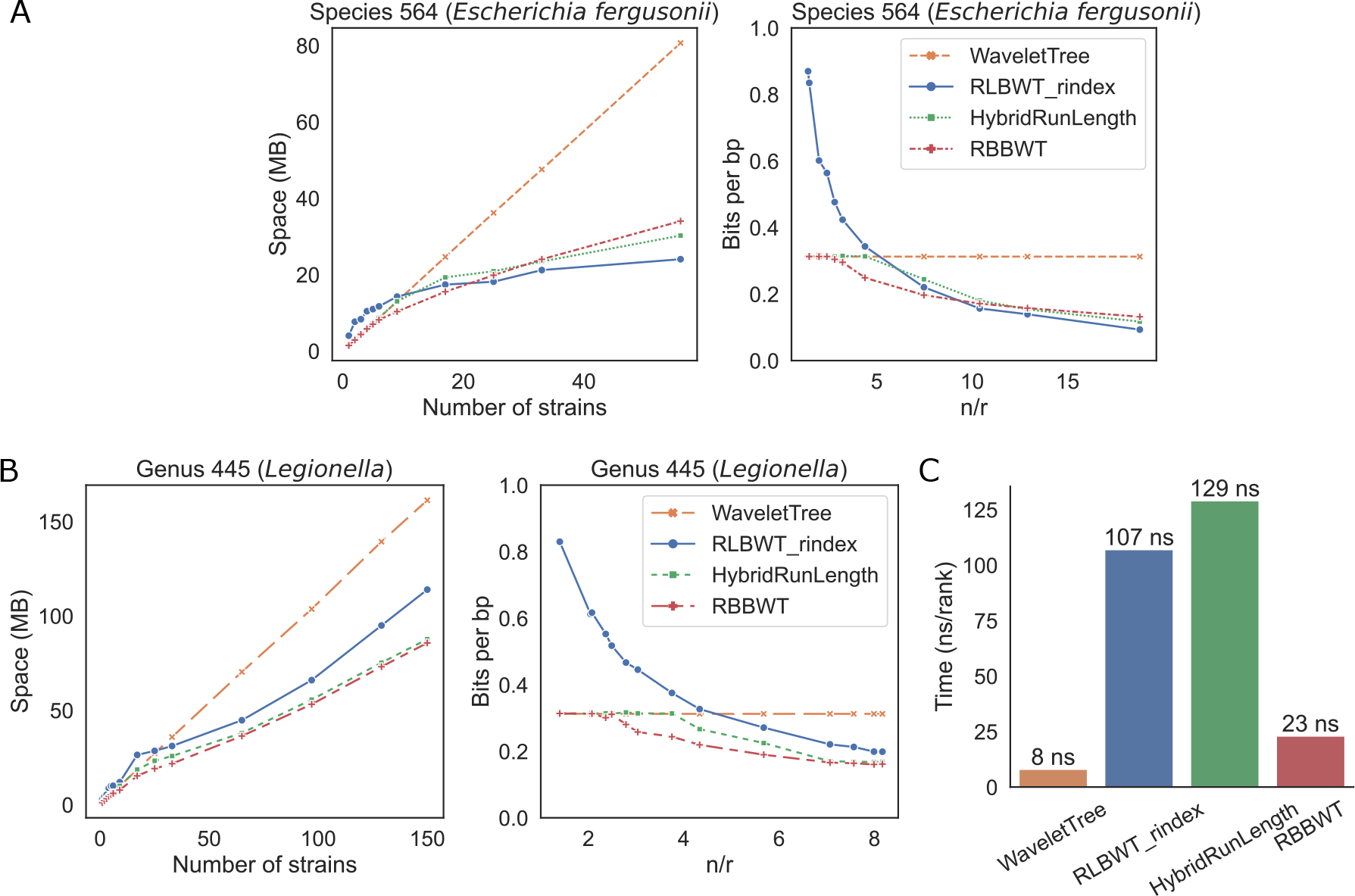
Computational efficiency of the wavelet tree, RLBWT, hybrid run-length compression and RBBWT. (A) Space usage when representing increasingly more genomes with species ID 564 (*Escherichia fergusonii*). Leb: absolute space usage in megabytes (MB). Right: bits used to represent one base pair (bp) when the average run length of the BWT sequence (n/r) increases. (B) The space usage when representing genomes with genus ID 445 (*Legionella*). Leb and right plots represent the same analysis as in (A). (C) Rank query time.

We next compared the space usage when compressing the BWT sequence for genomes from the same genus. We examined the genus *Legionella* (taxonomy ID 445) containing 150 genomes. Since genomes from the same genus were more diverse than genomes from the same species, the final L is 7.1 after adding all the non-plasmid sequences. RBBWT consumed the least memory among the benchmarked compression methods, using 46.9%, 24.8% and 2.3% less space than wavelet tree, RLBWT and hybrid run-length compressed BWT, respectively, in the end (Figure 2B). We further compared the speed of rank queries by averaging the time for finding the rank of ‘A’ for each of the first ten million positions in the final BWT, i.e., the average time of calling rank_’A’_(1, BWT) to rank_’A’_(10000000, BWT). Rank query in RBBWT was about five times faster than in RLBWT and only three times slower than using a wavelet tree (Figure 2C). Hybrid run-length compression was the slowest method. In both the species ID 564 and the genus ID 445 analysis, the block size of RBBWT was automatically determined to be 8 (Method), supporting the mild repetitiveness in the microbial genome database. To summarize, RBBWT is more memory-efficient and supports faster rank queries compared to RLBWT when compressing microbial genomes.

### Performance on classifying simulated data

We compared Centrifuger, Centrifuge and Kraken2’s accuracy on one million simulated paired-end short reads from 34,190 prokaryotic genomes (RefSeq bacteria+archaea). All three methods built the database indices on the same set of genomes. The average run length of the BWT was about 6.8, and the block size of RBBWT was automatically determined to be 8, the same value inferred as in the *Legionella* genome analysis above. We used TP (true positive) for the number of reads that are correctly classified at the specified taxonomy node or in its subtree, T (true) for the number of input reads, and P (positive) for the number of reads that were correctly classified at the specified taxonomy level or below. Therefore, we define the sensitivity as 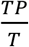, and precision as 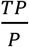. The strain-level classification evaluation in each method was for the reads classified to leaf nodes in the taxonomy tree. On this simulated data set, Centrifuger achieved the best accuracy at all taxonomy levels and was significantly better than Centrifuge and Kraken2 at species and genus levels (Figure 3A, Table S1). For example, Centrifuger’s sensitivity was 34.5% higher than both Centrifuge and Kraken2 at the species level. All three classifiers had comparable precision except at strain level. Centrifuge’s low sensitivity could be due to its policy of not resolving taxonomy IDs for matches hitting too many places in the database, where the threshold for the number of hits of a match was 40·report_threshold (default value of Centrifuge’s report threshold for the taxonomy IDs is 5). The strategy of handling matches on repetitive regions was one of the main differences between Centrifuger and Centrifuge during the classification stage (Method).

**Figure 3.**
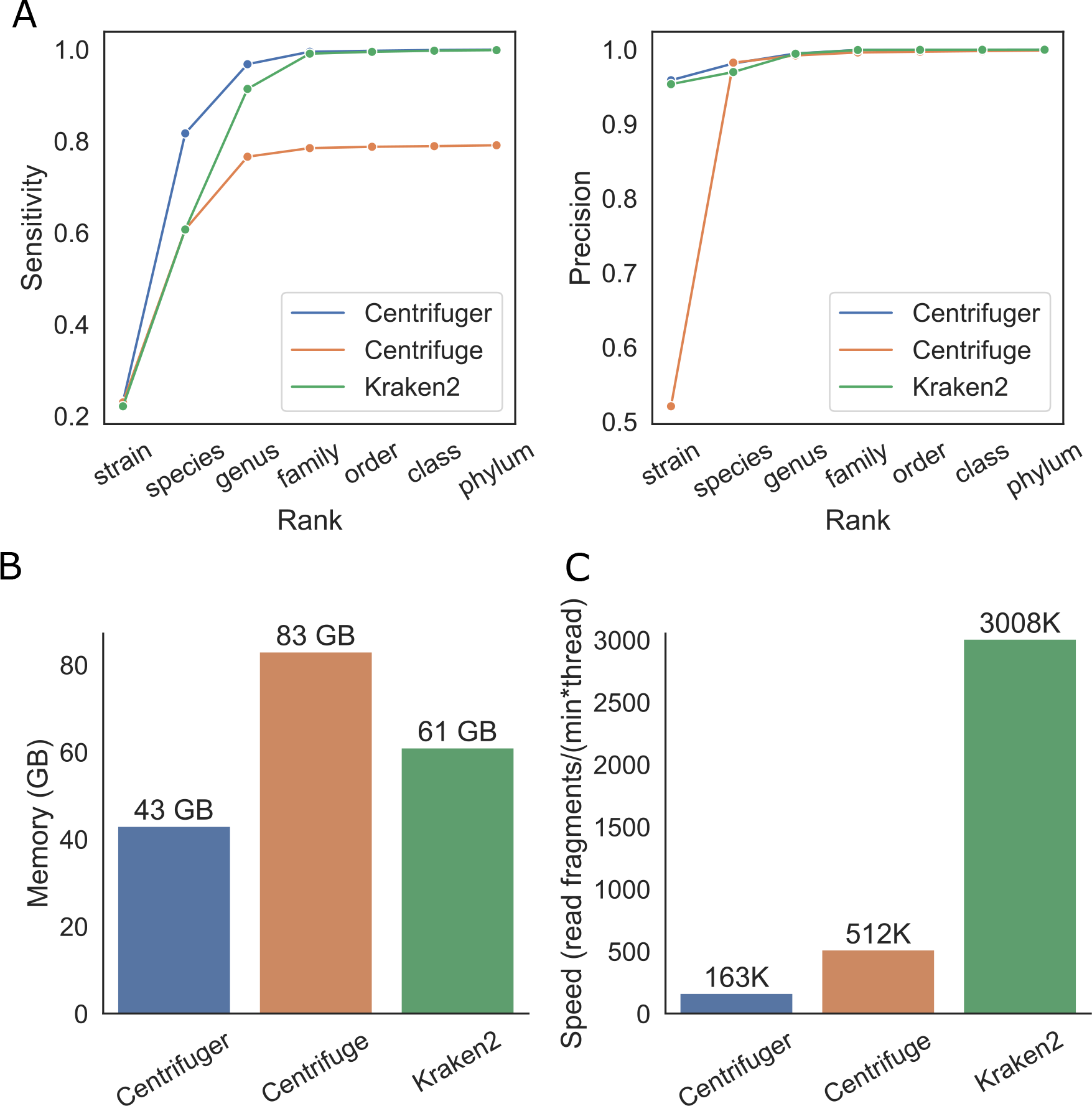
Performance of Centrifuger, Centrifuge and Kraken2 on the simulated data generated from June 2023 RefSeq prokaryotic genomes. (A) Sensitivity (leb) and precision (right) of Centrifuger, Centrifuge and Kraken2 at various taxonomy ranks. (B) Peak memory usage of each classifier. (C) Classification speed of each classifier with a single thread.

We utilized this simulated data set to compare the classifiers’ computational efficiency. Centrifuger was the most memory-efficient method, classifying the reads against the 140 billion base-pair (GBp) database using 43 gigabytes (GB) of memory (Figure 3B). For the classification speed, Kraken2 was the fastest method. Centrifuger and Centrifuge were also efficient, and processed more than 100,000 read pairs per minute using a single thread (Figure 3C). Speed differences were smaller when running with multiple threads. With eight threads, Centrifuger and Centrifuge could respectively process 1.2 million and 2.7 million reads in a minute, while Kraken2’s speed was 6.7 million reads/minute.

We also evaluated the accuracy of these three classifiers on 10 simulated short-read samples from the Critical Assessment of Metagenome Interpretation 2 (CAMI2) [29] challenge datasets. Since the truth table from CAMI2 was mostly at the species level, we skipped the strain-level comparison. As in the previous simulated data evaluation, Centrifuger achieved significantly higher sensitivity and precision at species and genus levels than Kraken2 and Centrifuge (Figure 4). For example, at the species level, the mean sensitivity of Centrifuger was 72.9% and 54.1% higher than Centrifuge’s and Kraken2’s, and the mean precision was 8.3% and 11.0% higher than Centrifuge’s and Kraken2’s, respectively.

**Figure 4.**
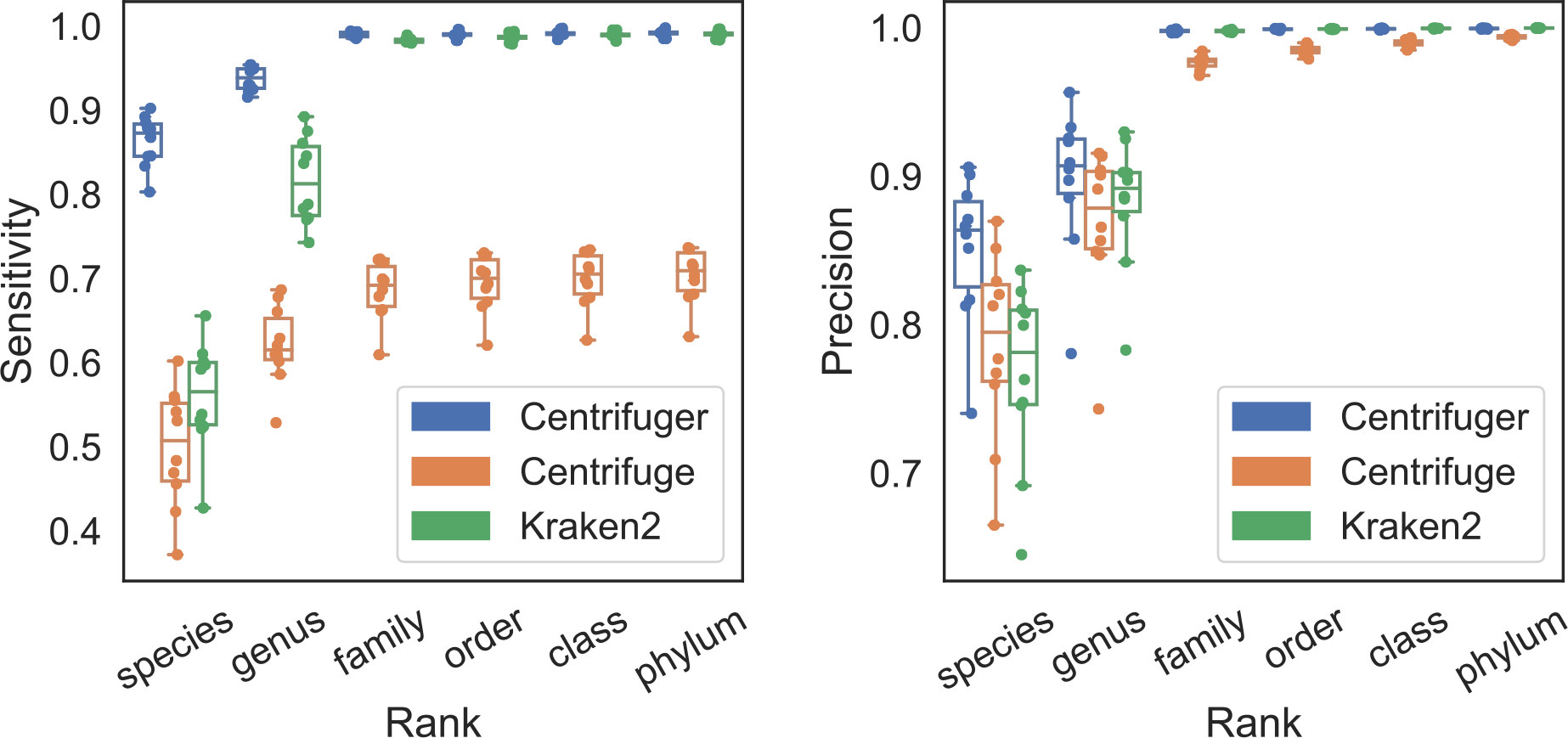
Sensitivity (leb) and precision (right) of Centrifuger, Centrifuge and Kraken2 at various taxonomy ranks on 10 simulated data sets from CAMI2

### Performance on classifying bacterial whole-genome sequencing data

We next evaluated Centrifuger, Centrifuge and Kraken2 on real bacterial whole-genome sequencing (WGS) data using the same database indexes as the simulated data evaluation. The true taxonomy IDs for each WGS sample were extracted in corresponding SRA RunInfo entries. We considered two scenarios: one where the RefSeq database contained some genomes from the same species (species-in), and one where the database did not include any same-species genomes but did include some same-genus genomes (species-not-in). Sensitivity and precision were defined in the same way as the simulated data evaluations, and we focused on the accuracy at the species and genus levels for species-in and species-not-in scenarios, respectively. For the species-in scenario, Centrifuger achieved 10.6% and 1.3% higher average sensitivity, 5.8% and 18.6% higher average precision than Centrifuge and Kraken2, respectively (Figure 5A). For the species-not-in scenario, while all three methods had comparable sensitivity, Kraken2’s precision was 10.2% and 21.2% higher than Centrifuger and Centrifuge, respectively (Figure 5B). Furthermore, species-not-in scenario has inferior accuracy compared with species-in scenario, suggesting that having a comprehensive database may substantially improve classification results by reducing the species-not-in chance.

**Figure 5.**
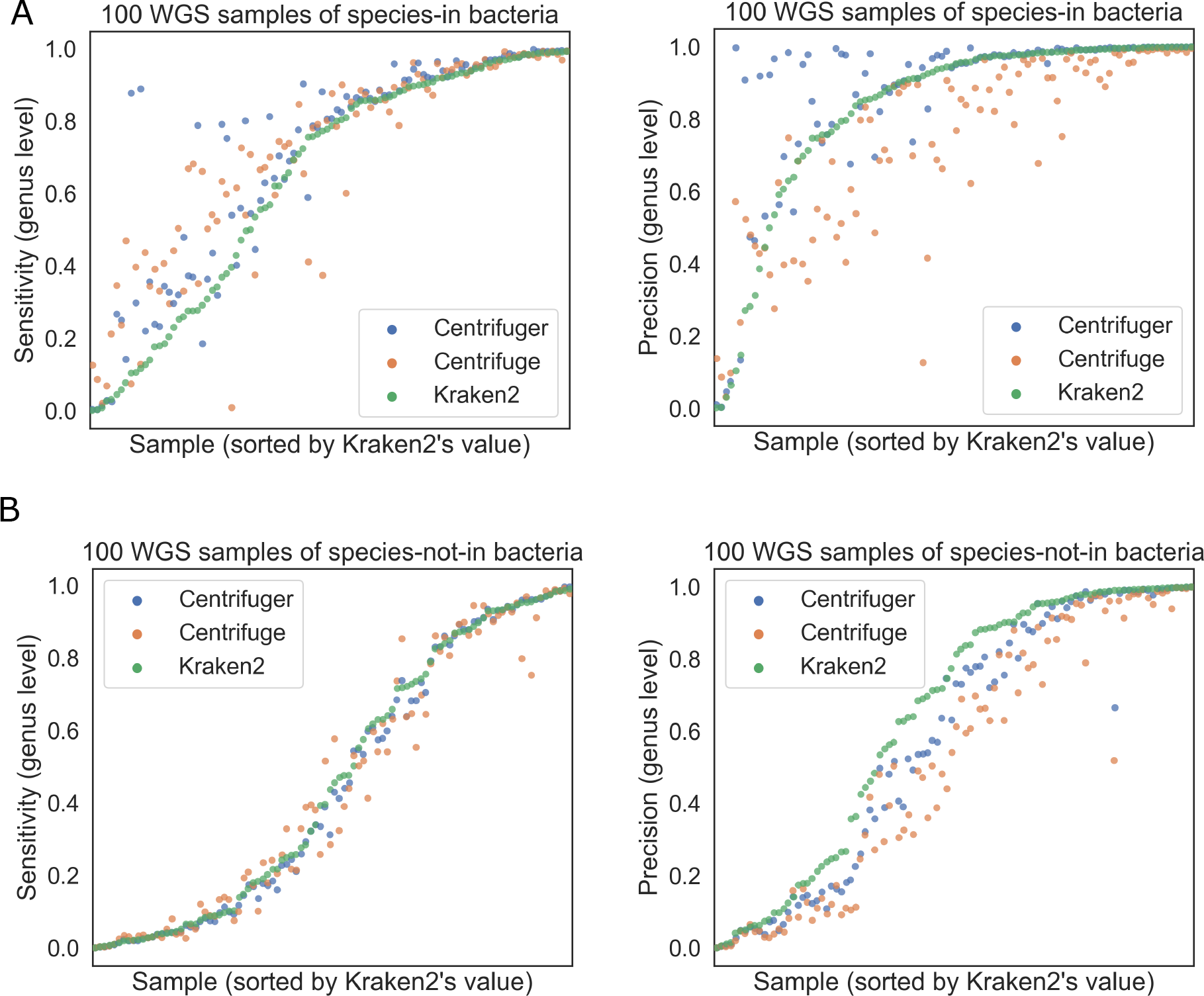
Performance of Centrifuger, Centrifuge and Kraken2 on bacterial WGS sequences (A) Sensitivity (leb) and precision (right) if species of bacteria are present in the database. (B) Sensitivity (leb) and precision (right) if species of the bacteria are not in the database but their genera are present in the database.

### Performance on classifying SARS-CoV-2 Oxford Nanopore WGS data

When a read can be uniquely classified to a sequence, Centrifuger reports the sequence ID in addition to the taxonomy ID, while Kraken2 provides only the taxonomy ID information. Centrifuger’s additional output is desirable for virus analysis. For example, SARS-CoV-2 variants’ genomes are all under the same taxonomy ID 2697049 in RefSeq and GenBank. To explore the effectiveness of sequence-level classification, we downloaded Oxford Nanopore WGS data from two SARS-CoV-2 projects with NCBI BioProject accession numbers PRJNA673096 and PRJEB40277, where PRJNA673986 were samples from US and PRJEB40227 were samples from Ireland. For each project, we selected 100 samples with the greatest number of reads (Table S2). Since RefSeq only had one SARS-CoV-2 sequence, we added the 92 SARS-CoV-2 sequences from GenBank and created an index comprising RefSeq human, prokaryotic, virus, and GenBank SARS-CoV-2 genomes. Centrifuger classified more than 99.9% of the reads to the taxonomy ID 2697049 on average, among which 24.1% were also identified with sequence IDs. Even though reads from a WGS sample should come from one variant, the true variant may not be present in the database, so reads can hit different variants. When examining the read fraction for each variant, namely sequence ID, the Irish samples and US samples showed distinct patterns (Figure 6A, heatmap with raw read fraction in Figure S1). We further conducted a principal component analysis (PCA) based on each sequence’s read fraction, normalized by the number of reads with sequence IDs. Samples from the US and Ireland were well separated into two clusters (Figure 6B) based on the first two principal components (PCs), suggesting that variants found in the two projects may have different sequence features. When inspecting the SAR-CoV-2 variant that contributed the most to the PC1 relative to the contribution to PC2, we found that variants detected in the Irish samples might have homologous regions to MT019531.1 while US samples did not (Figure 6C). On the other hand, when checking PC2’s major contributors, the MT158706.2 variant was commonly detected in both projects, suggesting that PC1 captured the project-specific variants.

**Figure 6.**
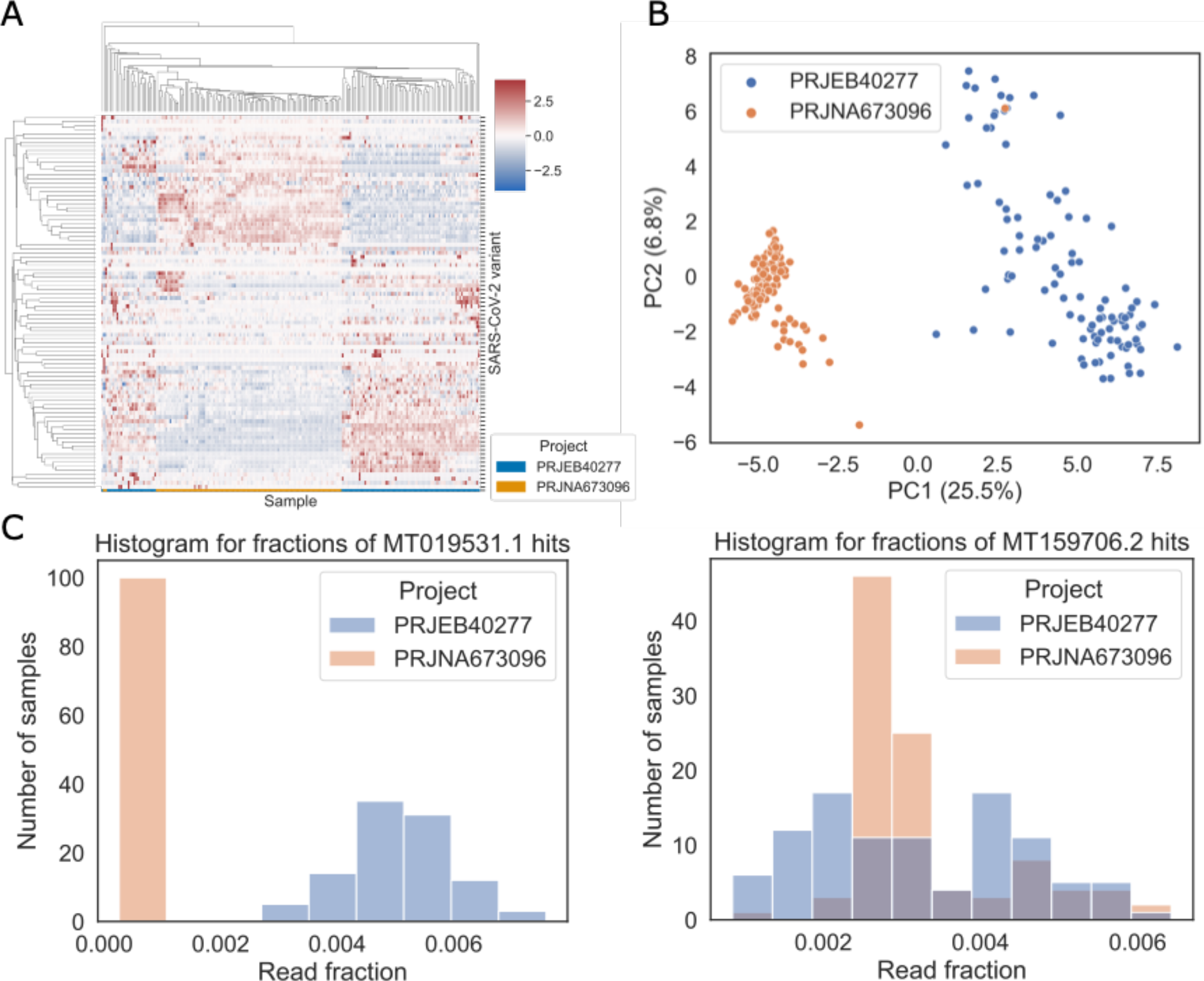
Sequence-level classification for SARS-CoV-2 WGS samples with Centrifuger (A)Fractions of reads hit on each SARS-CoV-2 variant. The rows are SARS-CoV-2 variants present in the RefSeq and GenBank, and the columns are the Oxford Nanopore WGS samples. The value for each row is standardized as z-scores. (B) PCA based on the read fraction for each variant. The numbers in the parenthesis are the variance fraction explained by each PC. (C) Leb: histogram of read fractions classified to MT01953.1 which has the most significant contribution to PC1 relative to its PC2’s contribution; right: histogram of read fractions classified to MT159706.2 which has the most significant contribution to PC2 relative to its PC1’s contribution.

## Discussion

We conducted comprehensive benchmarks to demonstrate that the RBBWT can significantly reduce the memory usage of the FM-index built over a microbial genome database. For additional space savings, we store the sequence ID rather than the full coordinate information in the sampled suffix array, a strategy also used in Centrifuge. The space for the sampled sequence IDs is further trimmed by bit-compact representation, making it a less impactful factor in space usage. For example, in Centrifuger’s 41-GB index of RefSeq prokaryotic genomes, 23 GB was for the RBBWT and 17 GB was for the sampled sequence IDs. Nevertheless, the overall space complexity of Centrifuger is still O(n) words due to the structure of sampled sequence IDs, making it worse than r-index’s O(r) words in the future as the repetitiveness of the genome database may grow fast. Though users can select a sparser sampling rate to maintain the space usage, this is at the expense of time efficiency. For instance, the index size became 32 GB when increasing the sampling rate from the default 16 to 32, but the classification speed of a thread decreased from 163K read/minute to 102K read/minute. Future work is needed to design a representation of the sampled sequence ID in sublinear space without sacrificing classification speed for microbial genomes.

Resolving the sequence IDs for each match is a time-consuming step in Centrifuger, especially when a match hits many sequences. Centrifuger’s current implementation follows the traditional FM-index paradigm, by repeatedly applying the LF mapping for each hit until reaching a sampled sequence ID. Alternative techniques like the document array profile [30] support rapid sequence ID retrieval, but are designed for highly repetitive genomes like human pangenome. Therefore, a memory-efficient algorithm for sequence ID resolving in microbial genomes is still needed. Since the default setting in Centrifuger is to report the LCA taxonomy ID for a read, algorithms like KATKA [31] that directly find the LCA taxonomy ID for a k-mer might suggest ways to avoid the overhead of repeated LF mapping in Centrifuger as well.

The current Centrifuger index stores nucleotide-based sequences. The Kraken2 and Kaiju [32] studies, however, observed that translated search, i.e., finding matches based on amino acids by translating nucleotides, could improve the classification accuracy for viral genomes. Since the wavelet tree data structure supports arbitrary alphabet sets, the RBBWT representation can be naturally extended to process amino acid sequences. RLBWT and RBBWT’s implementations are scalable for large alphabet set size, with a factor of log(σ) or the entropy in the space complexity. We found an alternative form of runlength encoded BWT, exported from ropeBWT2 [33] (with -dRo option) as the Fermi’s [34] format. RopeBWT2’s representation was 12.5% smaller than RLBWT on average with comparable rank query speed as RLBWT when compressing *Escherichia fergusonii*’s genomes. The slimmer size of ropeBWT2’s output might be attributed to its implementation being designed for small σ, where its space complexity is linear to σ. This may suggest that the efficiency of RBBWT and Centrifuger could be further improved for nucleotide search with tailored implementations that are less scalable for σ.

## Conclusions

Centrifuger is an efficient and accurate taxonomic classification method for processing sequencing data, including metagenomic sequencing data. Centrifuger adopts a novel compact data structure, run-block compressed sequence, to achieve sublinear storage space for BWT sequence without sacrificing much time efficiency. Specifically, Centrifuger can represent the 140 GBp Refseq prokaryotic genomes with an index of size 41 GB and classifies about 163K microbial reads every minute per thread. Furthermore, the lossless representation nature and the unconstrained pattern match length help Centrifuger achieve significantly better accuracy, in both sensitivity and precision, for classifications at the species or genus level. We expect that Centrifuger will contribute to microbiome studies by allowing the incorporation of the recent, more comprehensive microbial genome database. Centrifuger is a free open-source sobware released under the MIT license, and is available at https://github.com/mourisl/centrifuger.

## Methods

### Sequencing data and benchmark details

We generated the simulated data from the current RefSeq database using Mason v0.1.2 [35] with the option “illumina -pi 0 -pd 0 -pmms 2.5 -s 17 -N 2000000 -n 100 -mp -sq -hs 0 -hi 0 -i”, which simulated reads with 1% error rate. We further filtered the reads from the sequences without taxonomy information and kept the first one million read pairs as the final simulated data set. In addition to our own simulated data, we downloaded the first 10 simulated samples from the CAMI2 Challenge datasets at https://frl.publisso.de/data/frl:6425521/strain/short_read/. For the bacteria WGS data, we first downloaded the RunInfo from NIH NCBI SRA using the search word “(“Bacteria”[Organism] OR “Bacteria Latreille et al. 1825”[Organism]) AND (“2022/01/01”[MDAT] : “2023/08/01”[MDAT]) AND (“biomol dna”[Properties] AND “strategy wgs”[Properties] AND “plasorm illumina”[Properties] AND “filetype fastq”[Properties])”. Then based on whether the species or genus is present in the RefSeq database, we randomly pick 100 SRA IDs (Table S3), without repeating species or genus, for species-in and species-not-in evaluations, respectively.

We benchmarked the performance of Centrifuger v1.0.0, Centrifuge v1.0.4 and Kraken2 v2.1.3 in this study. Taxonomy information and microbial genomes were downloaded using the “centrifugerdownload” script in June 2023. Each classifier was used to build its own index on dustmasked [36] genome sequences. Kraken2 used its own built-in masking module. Centrifuger, Centrifuge and Kraken2 were tested with default parameters, except the additional “--no-abundance” option was specified for Centrifuge. We used the “centrifuge-promote” script in the Centrifuge package to reduce the classification result for each read to its LCA when calculating sensitivity and precision. All the benchmarks were conducted on the 2.8 GHz AMD EPYC 7543 32-Core Processor machine with 512 GB memory. The memory footprint was measured as the “Maximum resident set size” value from the “/usr/bin/time -v” command. When measuring speed, each classifier was run four times. The reported classification speed was calculated by taking the fastest runtime after excluding index loading time.

### Run-block compressed sequence

Run-block compressed sequence is a compact data structure supporting rank queries for any position in a sequence. For the input sequence T of length n and alphabet set Σ of size σ, we first partition T into equal-size substrings (blocks), *T*_1_, *T*_2_, …, *T*_*m*_, where 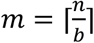 and *b* is the block size. The first component of the run-block compressed sequence is a bit vector *B*_*R*_ of size m indicating whether the corresponding block is a run block, i.e. a block consisting of one alphabet character repeated b times. We will then split T into two subrings, by concatenating run blocks and non-run blocks, i.e., 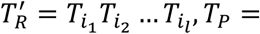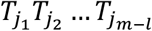, and 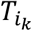 is the k-th run blocks in T with the alphabet 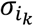. In the notation, we will use the subscript “R” to denote run-block compressed sequence, and “P” to represent the plain uncompressed sequence. Since the last block can still be determined as a run or non-run block even if it is shorter than *b*, we can assume that *n* is divisible by *b* for simplicity. The *T*_*R*_ can be lossless represented as 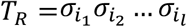, where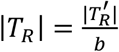 . The space-saving comes from using one character to represent a run block of size b, a strategy we call run-block compression. For example, for a sequence T=“TTTTACGTTTTTT”, when b=4, it will be split into “TTTTTTTT” and “ACGT” guided by *B*_*R*_ = 101. For the subsequence formed by the run blocks, we will use one character to represent each block in it. Therefore, the example sequence T will be represented by two sequences *T*_*R*_=“TT” and *T*_*P*_=“ACGT”, reducing the original length from 12 characters to 6 characters (Figure 1). We next show that run-block compression allows fast rank queries and sublinear space usage as the repetitiveness in the sequence increases. The rank query is the core operation in LF mapping during the backward search in FM-index.

#### Theorem 1.

The time complexity for rank query on run-block compressed sequence is *O*(*log*σ).

Proof: We will use the function *rank*_*c*_(*i, T*) to denote the rank for the alphabet c at position i of text T, where the index is 1-based. Equivalently, *rank*_*c*_(*i, T*) counts the number of c’s that occur before T[i], including T[i]. We can decompose the *rank*_*c*_(*i, T*) to the sum of corresponding ranks with respect to *T*_*R*_ and *T*_*p*_. There are two cases, depending on whether i is in a run block or not. Let k denote the block containing i, namely 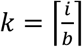. We compute the number of run blocks and non-run blocks before the block containing k as *r*_*r*_ = *rank*_1_(*k, B*_*R*_), a*n*d 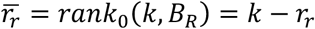, respectively. The *rank*_1_(*k, B*_*R*_) is the conventional rank query on bit vectors counting the number of 1s before or on position k in BR. With these notations, we can write the equations to compute *rank*_*c*_ (*i, T*).

When i is in a run block, i.e., *B*_*R*_ [*k*] = 1, we have:

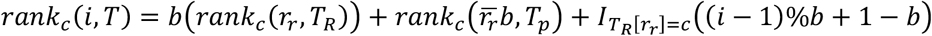

where 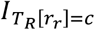 is an indicator that equals 1 if the subscript is true and equals to 0 otherwise. The last term is the special treatment if i is in a run block with alphabet character c. When i is in a non-run block, we have:

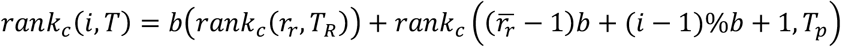

If we apply the wavelet tree to represent *T*_*R*_ and *T*_*P*_, then *rank*_*c*_(*i, T*) can be answered by at most two wavelet tree rank queries, one rank query on bit vector *B*_*r*_, and one wavelet tree access on *T*_*R*_.

Therefore, the total time complexity is 3*O*(*log*σ) + *O*(1) = *O*(*log*σ). □

The naïve implementation for calculating 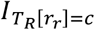 is to access the value of *T*_*R*_ [*r*_*r*_], requiring *O*(*log*σ) time if using wavelet tree. We note that the 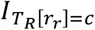 can also be inferred during the wavelet tree’s rank query on *T*_*R*_, by checking whether the bits labeling the relevant root-to-leaf path form the bit representation for c. This strategy further accelerates the *rank*_*c*_(*i, T*) operation, and is also applicable to other compressed sequence representations, including RLBWT.

#### Theorem 2.

The space complexity of run-block compressed sequence is 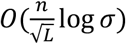 bits, where 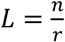 is the average run length and r is the number of runs in the sequence.

Proof: The key observation is that each non-run block contains at least one run head. Therefore, we have at most r non-run blocks. As a result, the minimum length of *T*_*R*_ is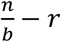, and the maximum length of *T*_*R*_ is *rb*. The length of the *B*_*R*_ is 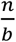 .

We can use wavelet trees to represent the run-block subsequence and the plain subsequence. Let *A* be the number of bits to represent one character in the wavelet tree, then the asymptotic total space usage in bit is 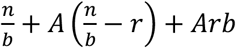, where the terms are for *B*_*R*_, *T*_*R*_, and *T*_*P*_, respectively. We can rewrite the space usage bound as:

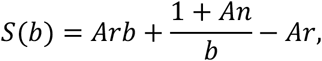

which is minimized when 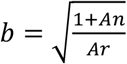. Because b is an integer, we take the block size as 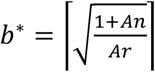.

Substituting this for *b*, we have:

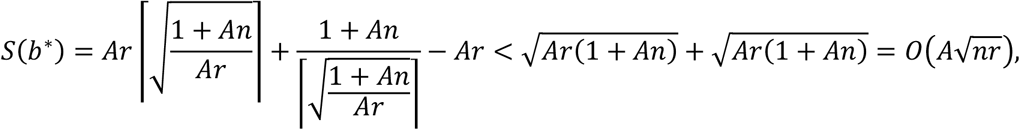

where the inequality is based on 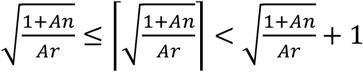. We can rewrite *S*(*b*^*^) by using the definition of 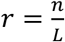, obtaining 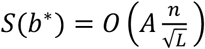. To ensure the worst rank query time on the data structure is small, the wavelet tree is in the shape of a balanced binary tree in our implementation, and *A* = *O*(logσ). Further space could be reduced if we use techniques like Huffman-shaped wavelet tree [37], and A will be *O*(*H*_0_(*T*_*R*_)) and *O*(*H*_0_(*T*_*P*_)) for T_R_ and T_P_, respectively, where H_0_ is the Shannon entropy of the sequence. □

The *b*^*^ found in the proof is to bound the worst-case space usage, where each non-run block has exactly one run head in the middle. The optimal block size can be different. For example, when every run has an identical length, the optimal block size is 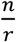 and every block is run-block compressible, yielding 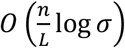-bit space complexity. To find the appropriate block size efficiently, we search the size of powers of 2s, e.g. 4, 8, 16,.., and select the block size 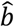 with the least space usage among them. Suppose 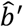 is the smallest power of 2 that is larger or equal to the block size *b*^*^ defined in the proof, then we have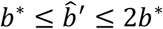. Therefore, 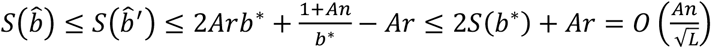, where the second inequality is by applying 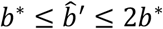. Therefore, the block size inferred from inspecting powers of 2 is not a bad estimator and gives the same asymptotic space usage as *b*^*^ in the worst case. Furthermore, since 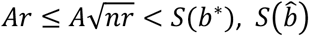 is no more than three times of the *S*(*b*^*^). To reduce the bias of the sparse search space, we also inspect the space usage of block sizes *b*^*^ and 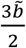 before making a final decision. When L is small, the block size minimizing the overall space usage is the length of the genome (TP=T), and run-block compression is equivalent to the wavelet tree representation. In practice, Centrifuger uses the first one million characters of the BWT sequence instead of the full sequence to infer the block size.

### Index construction

Centrifuger uses the blockwise suffix sorting algorithm [38] to build its index, as in Bowtie [39] and Centrifuge. The advantage of blockwise suffix sorting is to control the overall memory footprint and parallelize the construction procedure. The array that holds the BWT sequence is pre-allocated, and the sequence is filled in block-by-block as blocks of the suffix array are constructed. Another important component in the FM-index is the sampled suffix information. During construction, the index stores the genome coordinate information for every 16th offset, which can be adjusted by the user. Aber that, the offsets are transformed into strain-level sequence IDs and using a bit-efficient representation of the IDs.

For example, the RefSeq prokaryotic genome database contained 75,865 sequences (including plasmids) from 34,190 strains with complete genomes, so sequence IDs can be distinguished by a 17-bit integer.

Instead of saving the IDs in an array of 32-bit integers, the bit patterns are stored consecutively without any wasted space. Therefore, the total size for the bit-compact array for the sampled ID list is 17·m bits, or 0.26·m 64-bit words, where m is the number of sampled IDs.

### Taxonomic classification

For taxonomic classification, Centrifuger follows Centrifuge’s paradigm by greedily searching for semi-maximal exact matches in the database (Figure 1). For each read pair, Centrifuger searches the matches twice, one using the forward strand and the other using the reverse-complement strand. Let *l* denote the length of a match. Centrifuger will filter short matches where 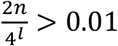 as they are likely from random hits. For example, we kept the matches with length greater or equal to 23 as valid matches when classifying reads using the 140 GBp RefSeq prokaryotic database. Each valid match will contribute a score (*l* − 15)^2^ to the corresponding strand, and the matches from the less-scored strand will be removed.

Centrifuger resolves sequence IDs contained in each valid match by using LF-mapping in the compressed FM-index to find sampled sequence IDs. If both strands have an equal score, the sequence ID will be resolved for both strands. For each resolved sequence ID, Centrifuger will sum up its scores across the matches using the formula *score*(*i*) = ∑_*i*∈*m*_(*l*_*m*_ − 15)^2^, where *i* ∈ *m* means match m hits the sequence with ID i. If a match spans a large range on the BWT sequence in the FM-index, i.e., it shows up many times in the database, Centrifuger will resolve the sequence IDs for at most 40·report_threshold (default report_threshold is 1 in Centrifuger) entries evenly distributed in the BWT interval. Though this is a heuristic that can cause the algorithm to miss the true genome of origin, it is likely to generate scores for sequences in the same phylogeny branch and may help identify the correct taxonomy IDs at higher levels. When the number of highest-scoring sequence IDs is more than the report threshold, Centrifuger will merge the IDs to their LCAs in the taxonomy tree until the number is within the threshold.

### Hybrid run-length compression

In addition to the run-block compression, we designed another compression scheme called hybrid runlength compression, using run-length compression for each fixed-size block. Hybrid run-length compressed BWT uses the same amount of space as the wavelet tree representation when the repetitiveness of the sequence is low, and its space usage converges to the RLBWT’s when the repetitiveness grows. In our implementation, a block is marked as run-length compressible (BR=1) if its average run-length is more than six based on the comparison between RLBWT and wavelet tree (Figure 2A,B). The substrings from blocks are separated into two subsequences based on BR, and the subsequences will be concatenated into two sequences, TR and Tp, respectively. TR will be compressed by the run-length method as in RLBWT, and Tp will be represented as a wavelet tree. The block size is inferred in the same fashion as in RBBWT. The rank query on the hybrid run-length compressed sequence is like the run-block compression, where we combine the ranks from TR and TP. This idea is similar to the wavelet tree when using fixed-block boosting [40]. Our implementation avoids explicitly recording the accumulated count at the beginning of a block and is therefore well suited to mildly repetitive sequences needing small blocks. For example, the block size of the hybrid run-length compressed BWT was only 12 for the genus *Legionella*’s genomes.

## Declarations

## Availability of data and materials

The source code of Centrifuger is available at https://github.com/mourisl/centrifuger. The code for the evaluations and experiments is available at https://github.com/mourisl/centrifuger_evaluations. The index for the RefSeq prokaryotic, human, virus and GenBank SARS-CoV-2 variants is available at Zenodo doi:10.5281/zenodo.1002323ti.

## Authors’ contributions

L.S. conceived the project. L.S. and B.L. designed the algorithm. L.S. implemented and evaluated the sobware. L.S. and B.L. wrote the manuscript.

## Abbreviations

FM-index: Ferragina-Manzini index
BWT: Burrows-Wheeler transform
RBBWT: run-block compressed BWT sequence
RLBWT: run-length compressed BWT sequence
CAMI2: Critical Assessment of Metagenome Interpretation 2
WGS: whole-genome sequencing
PC: principal components

## Acknowledgments

We thank Dr. Daehwan Kim and Dr. Florian Breitwieser for the foundation work in Centrifuge. We also thank Dr. Heng Li and Dr. Shannon Sousy for many helpful discussions.

---

**Table S1.** The classification accuracy at various taxonomy ranks in the Mason-generated simulated data. Sensitivity=TP/T, Precision=TP/P.

| rank | Matched (TP) | Predicted (P) | Cases (T) | sensitivity | precision | method |
| --- | --- | --- | --- | --- | --- | --- |
| strain | 231270 | 241184 | 1000000 | 0.2313 | 0.9589 | Centrifuger |
| species | 816975 | 832477 | 1000000 | 0.817 | 0.9814 | Centrifuger |
| genus | 967470 | 972537 | 999521 | 0.9679 | 0.9948 | Centrifuger |
| family | 992539 | 992922 | 997389 | 0.9951 | 0.9996 | Centrifuger |
| order | 995976 | 996205 | 998781 | 0.9972 | 0.9998 | Centrifuger |
| class | 996386 | 996466 | 997481 | 0.9989 | 0.9999 | Centrifuger |
| phylum | 999402 | 999448 | 999749 | 0.9997 | 1.0000 | Centrifuger |
| strain | 229849 | 441444 | 1000000 | 0.2298 | 0.5207 | Centrifuge |
| species | 607293 | 618127 | 1000000 | 0.6073 | 0.9825 | Centrifuge |
| genus | 765665 | 771680 | 999521 | 0.766 | 0.9922 | Centrifuge |
| family | 782725 | 785742 | 997389 | 0.7848 | 0.9962 | Centrifuge |
| order | 786747 | 788930 | 998781 | 0.7877 | 0.9972 | Centrifuge |
| class | 787116 | 788637 | 997481 | 0.7891 | 0.9981 | Centrifuge |
| phylum | 790843 | 791705 | 999749 | 0.791 | 0.9989 | Centrifuge |
| strain | 221359 | 232119 | 1000000 | 0.2214 | 0.9536 | Kraken2 |
| species | 607231 | 626061 | 1000000 | 0.6072 | 0.9699 | Kraken2 |
| genus | 913546 | 918606 | 999521 | 0.914 | 0.9945 | Kraken2 |
| family | 988327 | 988860 | 997389 | 0.9909 | 0.9995 | Kraken2 |
| order | 993684 | 994021 | 998781 | 0.9949 | 0.9997 | Kraken2 |
| class | 994895 | 995050 | 997481 | 0.9974 | 0.9998 | Kraken2 |
| phylum | 998390 | 998475 | 999749 | 0.9986 | 0.9999 | Kraken2 |

**Table S2.**
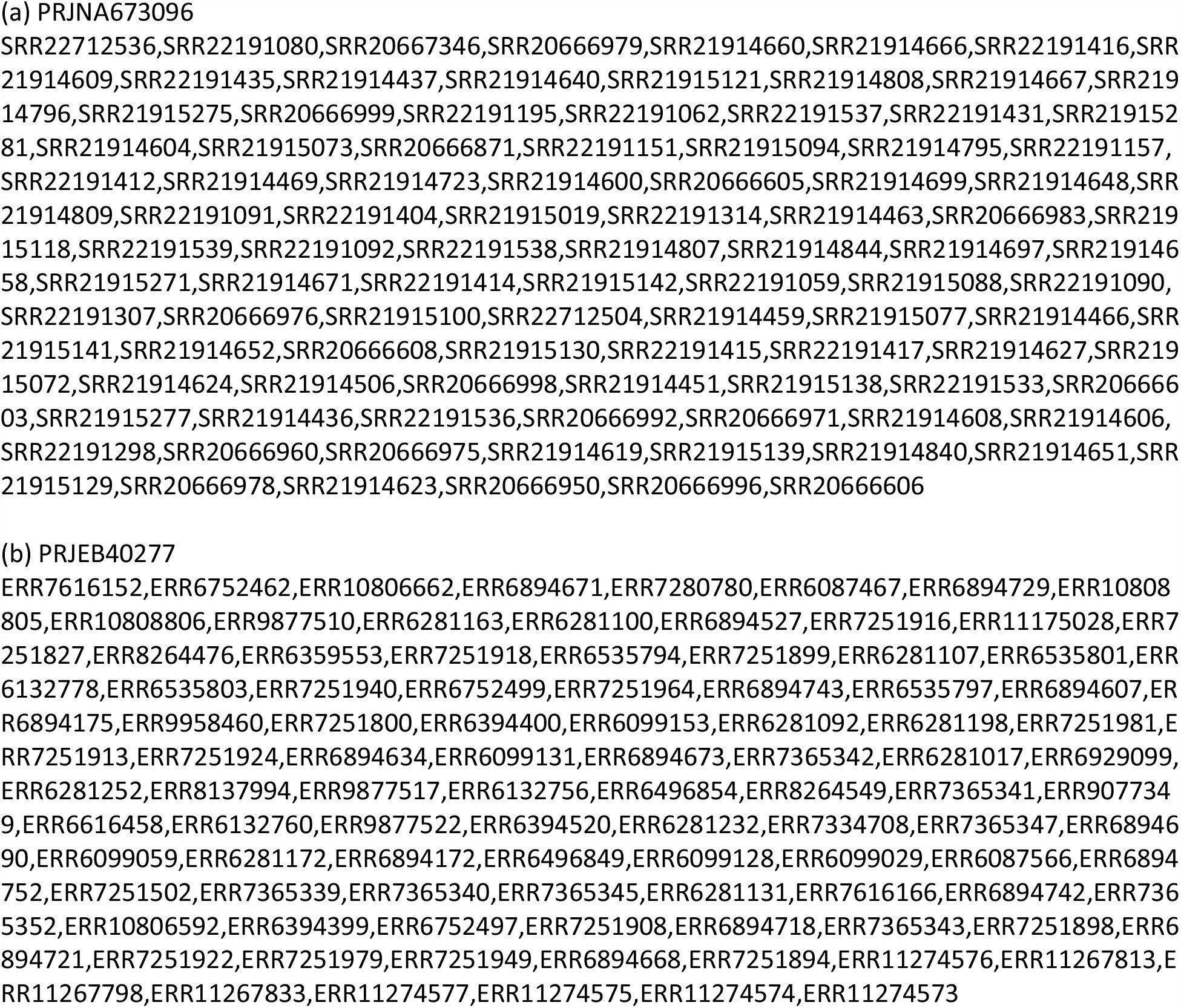
SRA IDs of the samples used in the SARS-CoV-2 sequence-level classification analysis.

**Table S3.**
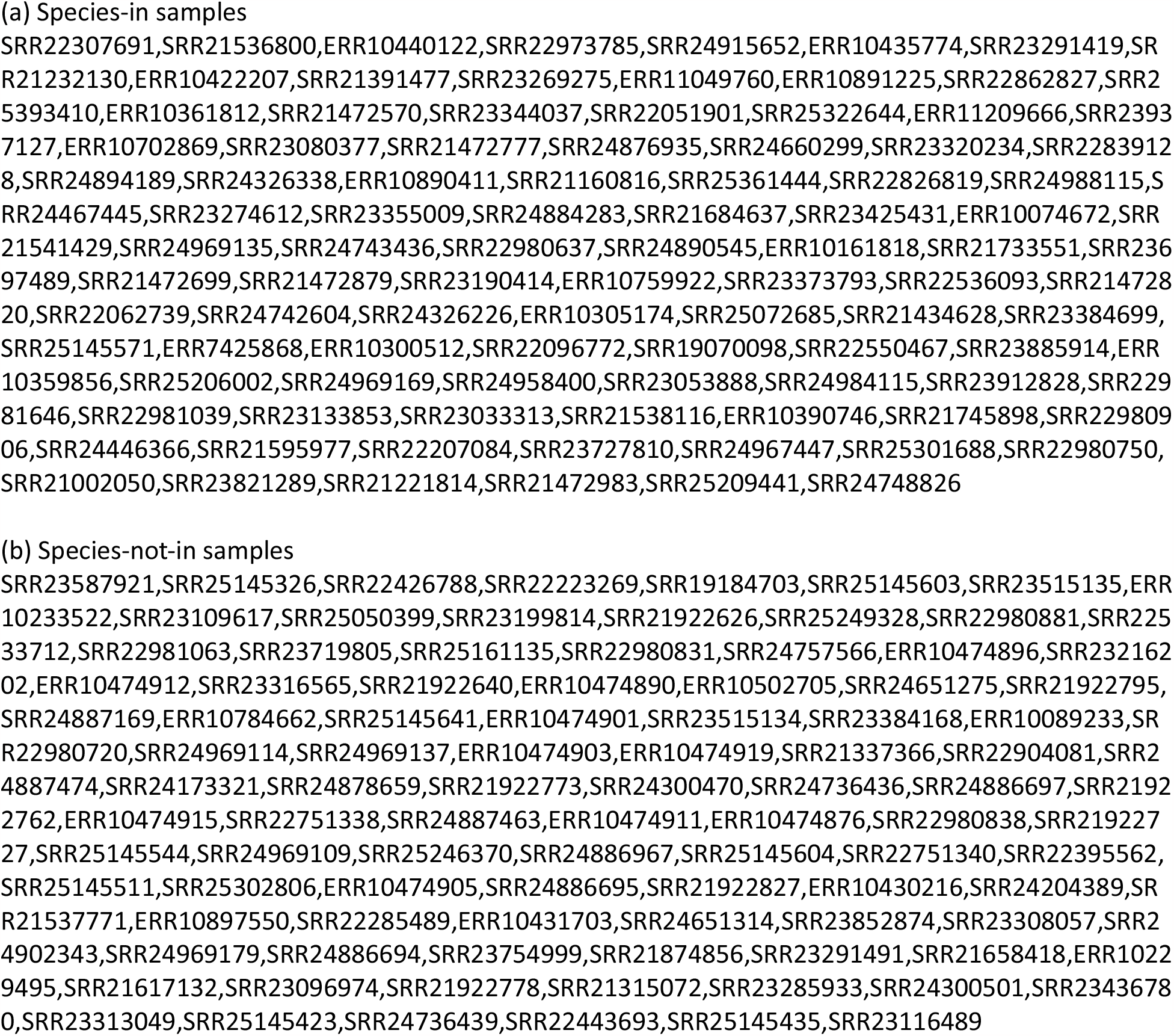
SRA IDs of the samples used in the bacterial WGS classification evaluations.

**Figure S1.**
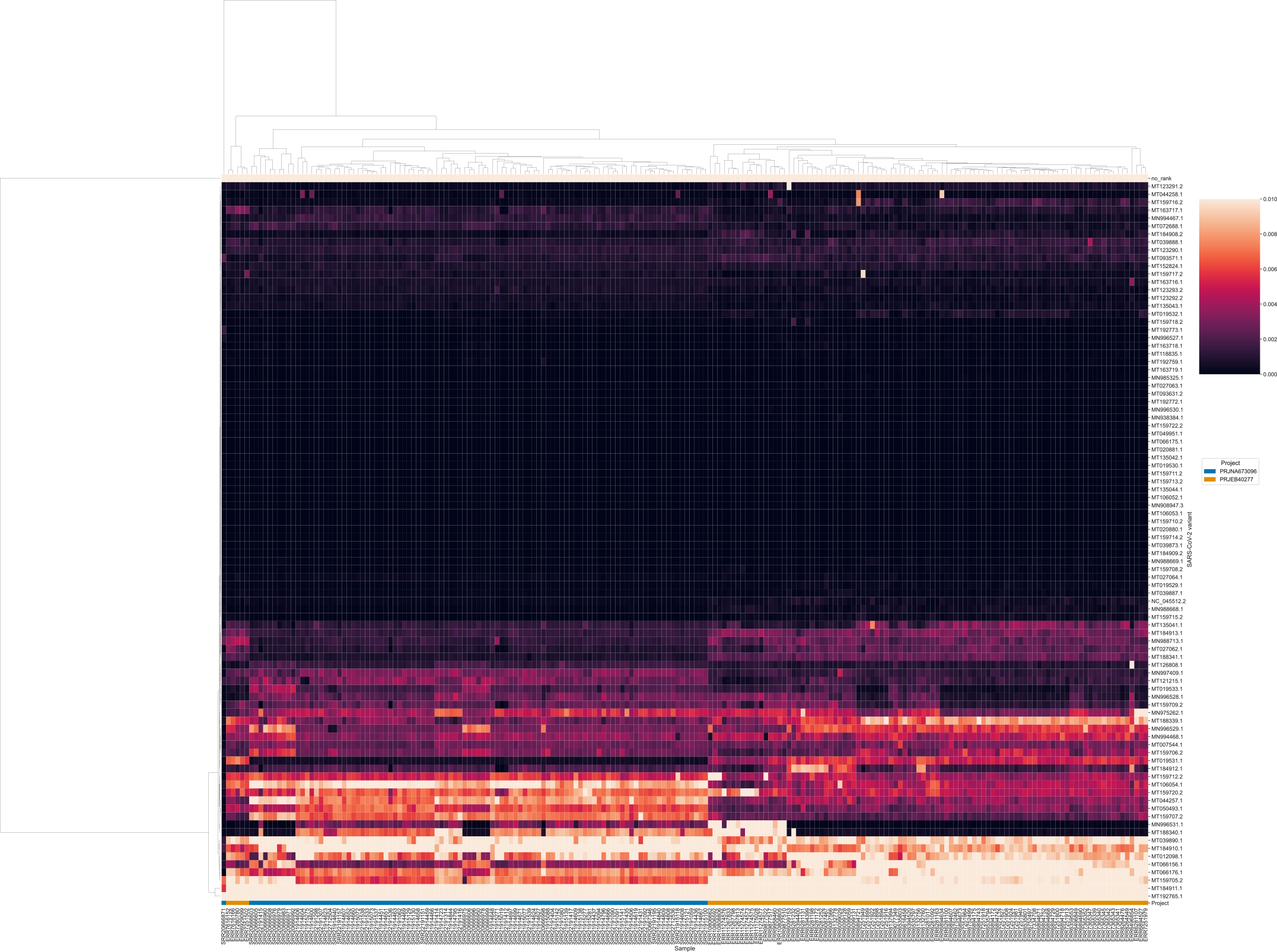
The heatmap with raw values of the read fractions for the SARS-CoV-2 sequence-level analysis

